# Preservation and changes in oscillatory dynamics across the cortical hierarchy

**DOI:** 10.1101/2020.02.03.932533

**Authors:** Mikael Lundqvist, André M. Bastos, Earl K. Miller

## Abstract

Theta (2-8 Hz), Alpha (8-12 Hz), beta (12-35 Hz) and gamma (>35 Hz) rhythms are ubiquitous in cortex. But there is little understanding of whether they have similar properties and functions in different cortical areas because they have rarely been compared across them. We record neuronal spikes and local field potentials simultaneously at several levels of the cortical hierarchy in monkeys. Theta, alpha, beta and gamma oscillations had similar relationships to spiking activity in visual, parietal and prefrontal cortex. However, the frequencies in all bands increased up the cortical hierarchy. These results suggest that these rhythms have similar functions inhibitory and excitatory across cortex. We discuss how the increase in frequencies up the cortical hierachy may help sculpt cortical flow and processing.

**Significance statement:** Phase-coupling in alpha/beta and gamma frequency ranges between cortical areas is often viewed as a means to shape brain-wide communication. However, systematic frequency differences between communicating areas are typically not considered, but equally important. Here we show that alpha/beta and gamma oscillations are of systematically higher frequency ascending the cortical hierarchy. This presents a fresh view on a widely studied topic. It has important implications in shaping cortical communication and helps explain widely observed phenomena.

## Introduction

Theta (2-8 Hz), alpha (8-12 Hz), beta (12-35 Hz) and gamma (>35 Hz) are common across cortex (Berger, 1929; Gray et al., 1990; Hari et al., 1997). Alpha/beta and gamma tend to be anticorrelated and have been associated with different functions. Increases in gamma power occur during sensory inputs/motor outputs while increases in alpha/beta have occur during top-down control and inhibition (Gray et al., 1990; Bastos et al., 2015, Buschmann and Miller, 2007; Van Ede et al., 2011, Lundqvist et al., 2016, Van Kerkoele et al., 2014, Fisch et al., 2009; Jokisch and Jensen, 2007; Pfurtscheller and Aranibar, 1977). For example, in motor cortex, beta power is high and gamma low when movements are inhibited but this reverses when movement is released (Pfurtscheller et al., 1996; Brovelli et al., 2004; Cheyne et al., 2008; Schmidt et al., 2019). In visual cortex, gamma power is high and alpha low during sensory stimulation; vice-versa for representations outside focal attention (Fries et al., 2001, Gray et al., 1990; Gevins et al., 1997; Fisch et al., 2009; Klimesch et al., 1998; Bollimunta et al., 2011; Buffalo et al., 2011; Pfurtscheller and Aranibar, 1977; Jokisch and Jensen, 2007; van Ede et al., 2011; Rohenkohl and Nobre, 2011). In the prefrontal cortex (PFC) bursts of gamma are associated with sensory information maintained in working memory (Lundqvist et al., 2016; 2018; Bastos et al., 2018). They are anti-corelated with alpha/beta bursts that decrease during encoding of sensory information in working memory and increase when working memory is cleared.

Given the ubiquity of these rhythms and their apparent push-pull relationship, it would important to know how they compare across cortical areas. Any preserved attributes of the rhythms would suggest shared roles in cortical processing while differences can provide insights into any functional differences across the cortical hierarchy. There is some evidence for differences. The lower rhythms skew towards alpha/low-beta in sensory and parietal cortex and towards higher beta in motor and prefrontal cortex, albeit in comparisons across different studies and species (Buschmann and Miller, 2007; Lundqvist et al, 2016; Bollimunta et al., 2011; Jokisch and Jensen et al., 2007). These differences could therefore be due to differences in task, species, and/or recording techniques. Few studies have compared multiple levels of the cortical hierachy using multiple intracranial electrodes that allow good localization of local field potentials as well as their relationship to neuronal spiking.

We recorded spikes and LFPs simultaeously along the cortical visual hierarchy as monkeys performed a visual working memory task. We analyzed local field-potentials and multi-unit activity. This revealed similar functional relationship between spiking, alpha/beta and gamma power across cortical areas. Alpha/beta power was lower during encoding and retention of information, and anti-correlated with spiking activity and gamma across time and recording sites. Both types of rhythms occurred in bursts, not sustained increases in power. Gamma was associated with spiking carrying sensory information; alpha/beta was negatively correlated with spiking. The peak frequencies of both rhythms increased as the cortical hierarchy was ascended from sensory to prefrontal cortex. Moreover, theta (2-8 Hz) often couples with gamma rhythms (Canolty et al., 2006; Schroeder and Lakatos, 2009; Tort et al., 2009). We found theta power was most pronounced in the higher areas, correlated positively with spiking and was also of gradually higher frequency ascending the cortical hierarchy. This suggets that interactions between alpha/beta and gamma are preserved across cortex and may play a general role in regulating expression of information by cortical neurons. We discuss the computational implications of the gradual increase of these frequencies up the cortical hierarchy.

## Materials and Methods

### Experimental design

Using positive reinforcement, we trained monkeys to perform a visual search task with a memory delay (Figure 1A). Monkeys fixated a point at the center of the screen (fixation window radius: 2-3 visual degrees) for a duration of 1 s. Then, one of three sample objects was presented at the center of gaze for 1 s. Then monkeys maintained central fixation over a delay (between 0.5-1.2 s, in some sessions (47/81) held fixed at 1.2 s). A search array then appeared. It consisted of an object that matched the sample together with either one or two distractor objects presented at the same eccentricity (3-8 degrees) but in different visual quadrants as the sample. The position of the match and the distractors were always randomly chosen. Monkeys were rewarded if they made a direct saccade to the match. Monkeys were trained on this task using a library of 22 sample objects. For recordings, we used a subset of these objects (12), choosing a total of 3 per session. The monkeys performed the task with blocks in which the sample was either randomly chosen trial-by-trial or held fixed. Only the data with trial-by-trial cuing, requiring engagement of working memory, were used and only the 81 sessions with at least 70 correctly performed trials included.

**Figure 1.**
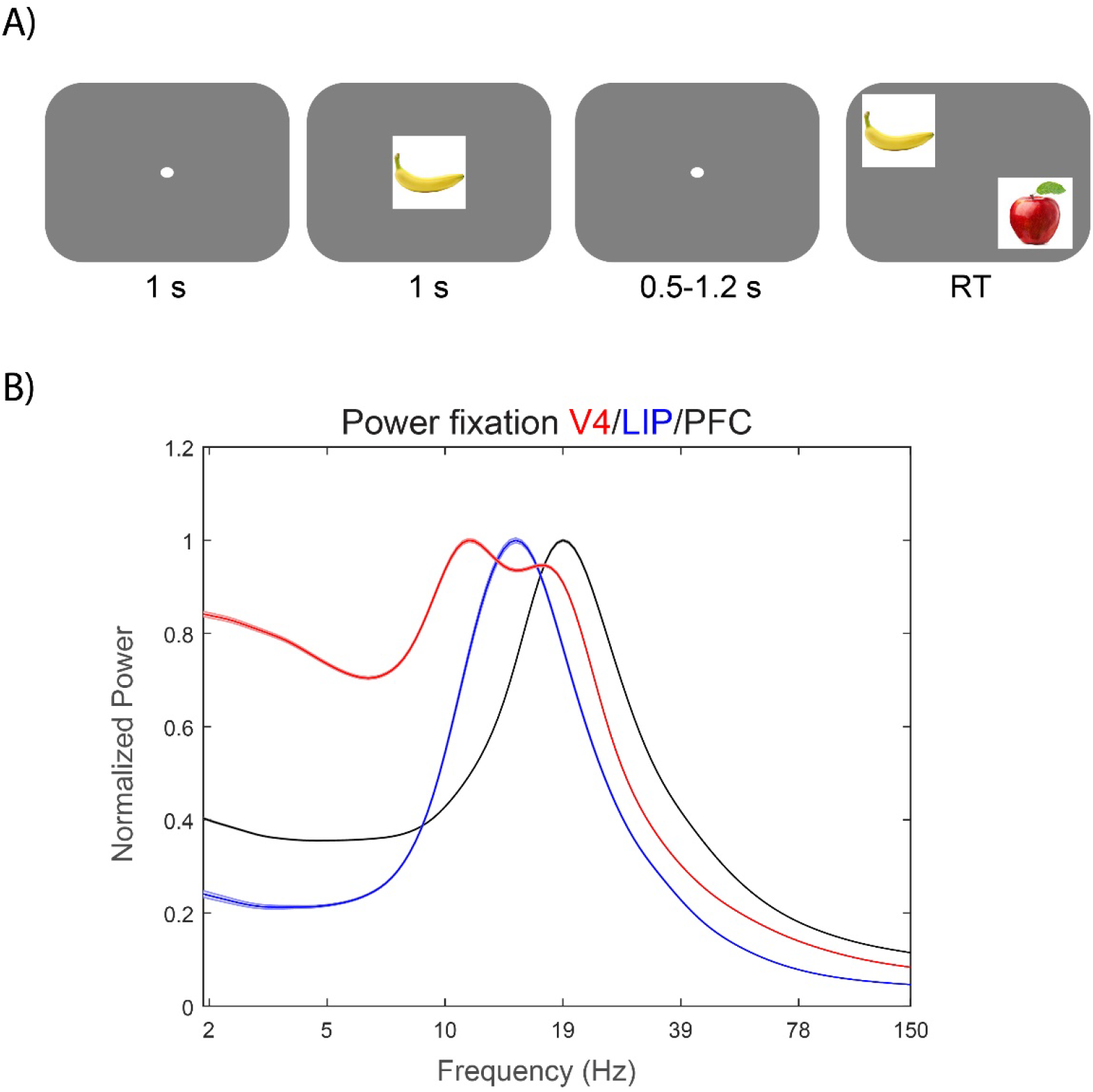
Task-structure and baseline power. A) Animals where to fixate at the center of the screen up until the response. A memory cue was presented foveally followed by a memory delay (0.5-1.2s). After the delay 2-3 items were presented peripherally and the animals where to saccade to the item that had been presented as a memory cue. B) Power spectrum from the middle of the fixation period (600-300 ms before stimulus onset) from V4 (red), LIP (blue) and PFC (black). Shaded areas show standard error of mean.

### Animal models

Two adult rhesus macaques (macaca mulatta) were used in this study (Monkey S: 6 years old, 5.0 kg and monkey L: 17 years old, 10.5 kg). Both animals were pair-housed on 12-hr day/night cycles and maintained in a temperature-controlled environment (80°F). All procedures were approved by the (Massaschsetts Institute of Technology (MIT) IACUC and followed the guidelines of the MIT Animal Care and Use Committee and the US National Institutes of Health.

### Data Collection

All of the data were recorded through Blackrock headstages (Blackrock Cereplex M, Salt Lake City, UT), sampled at 30 kHz, band-passed between 0.3 Hz and 7.5 kHz, and digitized at a 16-bit, 250 nV/bit. All LFPs were recorded with a low-pass 250 Hz Butterworth filter, sampled at 1 kHz, and AC-coupled.

We implanted the monkeys with three recording wells placed over visual/temporal, parietal, and frontal cortex. We kept a total of 64 sessions with multi-contact (“U probes” and “V probes” from Plexon, Dallas, TX) probes, and 17 sessions with acute single tip electrodes. In each session using multi-contact probes, we inserted between 1-3 laminar probes into each recording chamber with either 100 or 200 um intersite spacing and either 16 or 32 total electrodes per probe. Between 3-7 probes in total per session were used, with a total channel count ranging between 48-128 electrodes per session. The recording reference was the reinforcement tube, which made metallic contact with the entire length of the probe (total probe length from connector to tip was 70mm).

We used a number of physiologic indicators to guide our electrode placement, as previously described (Bastos et al., 2018). First, the presence of a slow 1-2 Hz signal, a heartbeat artifact, was often found as we pierced the pia mater and just as we entered the gray matter. Second, as the first contacts of the electrode entered the gray matter, the magnitude of the local field potential increased, and single units and/or neural hash became apparent, both audibly and visually with spikes appearing in the online spike threshold crossing. Once the tip of the electrode transitioned into the gray matter, electrodes were lowered slowly an additional ~2.5mm. At this point, we retracted the probe by 200-400 um, and allowed the probe to settle for between one to two hours before beginning the task. We left between 1-3 contacts out of gray matter in the overlying Cerebral Spinal Fluid (CSF). These contacts were not used in the analysis.

### Pre-processing

For the analysis of the analog multi-unit activity (MUA) we band-pass filtered the raw, unfiltered, 30kHz sampled data into a wide band between 500-5,000Hz, the power range dominated by spikes. The signal was then low-pass filtered at 250Hz and re-sampled to 1,000 kHz. The advantage of this signal is that it captures all nearby units, including those with low signal to noise ratio that would not be captured with a strict threshold. A subset of contacts had apparent artifacts. These contacts were automatically removed (16/768 in LIP, 12/2252 in PFC and 120/1538 in V4 (most from the same session)) from the analysis by setting a threshhold (100% above mean) on the power between 1-5 Hz.

### Analysis and statistical tests

All analysis was performed using Matlab. We estimated power at all frequencies from 2-150 Hz using Morlet wavelets (5 cycles, estimated each ms), with 81 frequencies (6 octaves) of interest distributed on a logarithmic scale.

We correlated MUA and power using neighbouring contacts to avoid spike bleed-through to drive correlations in the gamma band. Correlations were calculated for each frequency of interest, and we used Bonferoni correction to take this into account when calculating statistics. This was also the case for multiple pairwise t-tests between area pairs.

To determine stimulus selectivity we used bias-corrected Percentage explained Variance (PEV; Olejnik and Algina, 2003; Lundqvist et al., 2016). MUA was smoothed using 100 ms rectangular windows. For each 1 ms timepoint, a one-way ANOVA test was performed with trials grouped based on the sample cue identity. Non-biased PEV was used in order to avoid non-zero means for small sampe sizes.

We correlated the time-course of trial-averaged signals (power vs MUA or power vs PEV in MUA) for each electrode. For correlations over time we used the last 500 ms of fixation, the 1000 ms of sample presentation and the first 700 ms of the delay (not using trials with less than 700 ms delays). We also correlated, across electrodes, the change in MUA to the change in power from one epoch to another to determine relationships at the single electrode level. For this we used Spearman’s ranked correlations using the epoch averages per electrode. For power modulation by task (Figure 2) −700 to −300 ms was used for pre-stimulation and 200-1000 ms for stimulation, where 0 ms is the time of stimulation onset, to avoid frequency smoothing crosstalk between the epochs and frequencies of interest.

**Figure 2.**
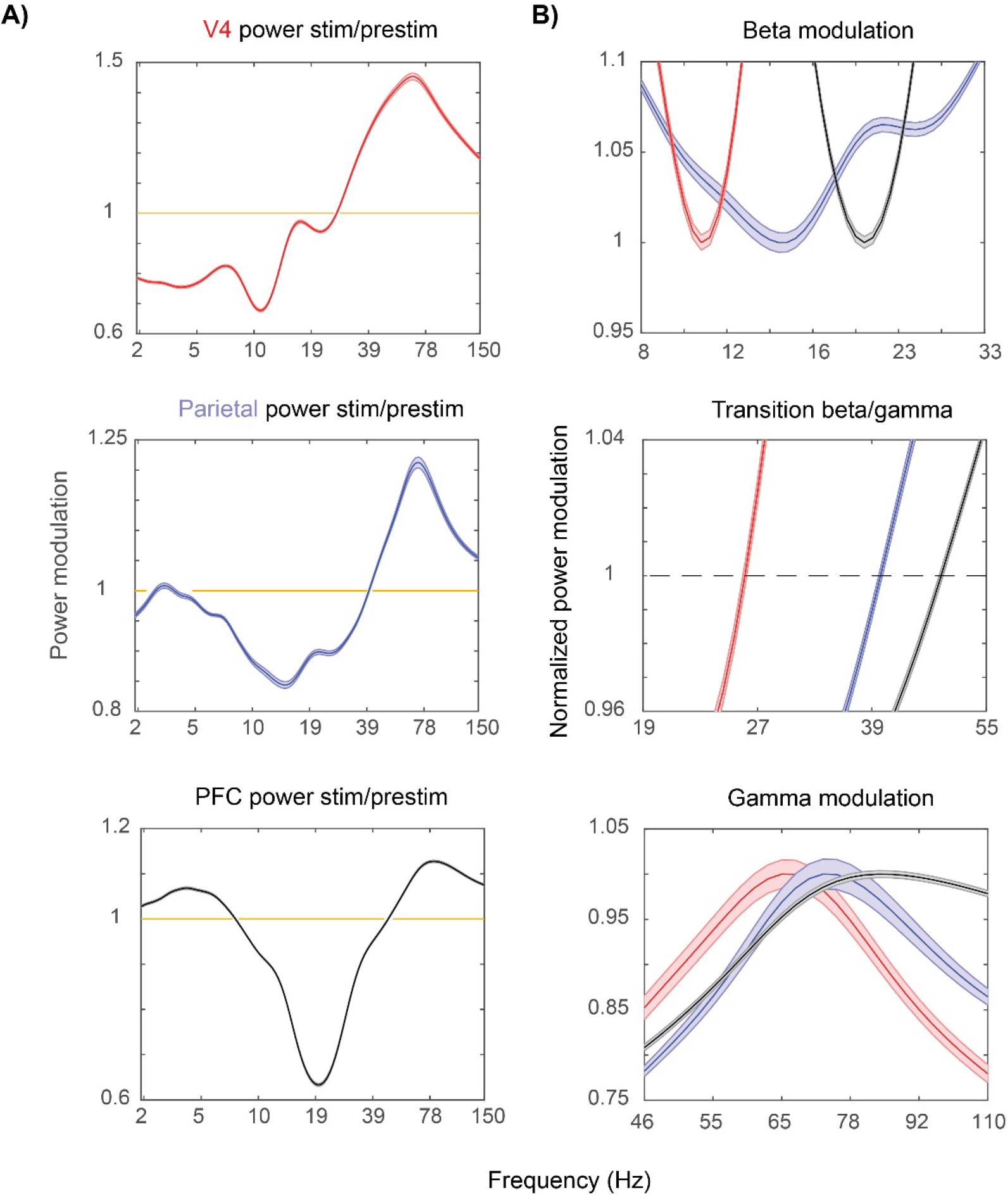
Stimulus induced power. A) show stimulus induced power (stimulus/pre-stimulus power; Methods) for V4 (top, red), LIP (middle, blue) and PFC (bottom, black). B) Zoom in on the beta suppression (top), transition from beta to gamma (middle) and gamma peaks (bottom) for V4 (red), LIP (blue) and PFC (black). Note that the peaks and beta throughs are normalized to 1 for easier comparison between areas in the plots in B). Shaded areas show standard error of mean.

In order to find spike-power relationships that did not depend on shared epoch preferences between the two measures we calculated correlations between neighbouring pairs of MUA and power and compared it to the correlations between random pairs of MUA and power (taken across all sessions but within each area). The correlations between random pairs were carried out 1,000 times to estimate a distribution that was then compared to the original correlations.

### Burst detection

Power displayed strong fluctuations on a single trial level, only exceeding baseline levels during brief burst events. To extract these we used a procedure developed earlier (Lundqvist et al., 2016). In short, we used the fixation period as baseline to calculate mean and standard deviation in each band of interest (beta, 6-35 Hz; gamma 1, 40-65 Hz; gamma 2, 55-90 Hz; gamma 3, 70-100 Hz). We used a sliding window of 10 trials to calculate the mean and standard deviation used for each trial. In a first step, candidate bursts were extracted if the average power within a given band exceeded the threshold (set at 2 standard deviations above the mean 10 trial average) for at least 2 cycles (based on the central frequency of each band).

Further, we extracted the time-frequency representation of the signal in the spectrotemporal neighborhood of each burst using the wavelets. We resorted to fitting two-dimensional Gaussian function to the local time-frequency map to specify the aforementioned neighborhood. Finally, we defined the burst length as a time subinterval where the average instantaneous power was higher than half of the local maximum (half-power point). The burst intervals were extracted for the alpha/beta band (6–35 Hz) and three gamma sub-band oscillations (40–65, 55–90, and 70–100 Hz) from each trial, along with the central frequency of each burst and the frequency width. The frequency width of bursts were estimated analogously to their duration: the frequency range where the spectral power component did not fall below 50% of the local maximum (epicenter of the burst’s power). We could then select bursts within a more narrow range than described by the original bands. The central frequency of each burst was used when correlating duration of bursts with their frequency. To determine burst durations of specific bands we only included bursts with their central frequency within a certain range of the average peak for that frequency band and area. For alpha/beta we used +−4 Hz and for gamma +−10 Hz. We used bursts from the first 70 trials for each session to get non-biased estimate.

### Data availability

The data and code will be made available by reasonable request by contacting the corresponding author, Earl Miller.

## Results

### Increase of frequencies up the cortical hierarchy

Two rhesus macaques performed a delayed-match-to-sample (DMTS) task (Figure 1A). We simultanously recorded local field-potentials (LFPs) and multi-unit activity (MUA) from V4 (n=1418 recording sites), Lateral intraparietal cortex (LIP, n=752) and prefrontal cortex (PFC, n=2240).

The mean peak frequency of oscillatory power increased up the cortical hierarchy. Figure 1B shows the average LFP power spectrum of each cortical area during a baseline interval just before presentation of the sample stimulus (see Methods). Each area was dominated by oscillations in the alpha (8-12 Hz) and beta (12-35 Hz) range (Figure 1B). The lack of a gamma peak was due to the lack of bottom-up sensory inputs, which are associated with increases in gamma power (Gray et al., 1990). The peak frequency of alpha/beta increased from V4 (11 Hz) to LIP (15 Hz) to PFC (19 Hz) (all across-areas comparisons had significantly different means at p<10^−30^, unpaired t-test corrected for multiple tests, using frequency between 7 to 40 Hz with maximum amplitude as peak at each recording site).

The modulation of oscillatory power by stimulus presentation also increased in peak frequency up the cortical hierarchy. We established the “functionally defined” frequency bands in each area by determing how each frequenciy was modulated by presentation of the sample stimulus. To do so, power per frequency during sample presentation was divided by the power per frequency during the baseline. The result showed that sample presentation suppressed alpha/beta power and increased gamma power (Figure 2). Note that the frequency of maximum suppression of alpha/beta matched the baseline peak frequency for each area. As such, it also increased from V4 (11 Hz) to LIP (16 Hz) to PFC (20 Hz) (all across-areas comparisons had significantly different means at p<10^−15^, unpaired t-test corrected for multiple tests on frequency of maximal decrease per electrode). The peak of average gamma power also increased in frequency from V4 (65 Hz) to LIP (72 Hz) to PFC (80 Hz)(Figure 2). We defined the transition between beta and gamma as the first frequency with positive modulation by stimulus presentation. This the transition also occurred at gradually higher frequencies as the cortical hierarchy was ascended (Figure 2B middle; V4: 26 Hz, LIP: 40 Hz, PFC: 47 Hz, all across-areas comparisons had significantly different means at p<10^−15^ using unpaired t-test corrected for multiple tests).

### Spiking was correlated with changes in oscillatory power

Increases in spiking were associated with increases in gamma and theta power and decreases in alpha/beta power. We examined the correlation between changes in MUA and oscillatory power across time (over three task epochs: baseline, sample presentation and memory delay). The level of MUA was positively correlated with changes in functionally defined (i.e., stimulus modulated, see above) gamma power and anti-correlated with changes in functionally defined alpha/beta power (Figure 3A; note that MUA and LFPs from neighboring, not the same, electrodes were used to avoid bleed-through effects). In LIP and PFC, but not V4, there was also a positive correlation between changes in theta power and level of MUA (Figure 3A).

**Figure 3.**
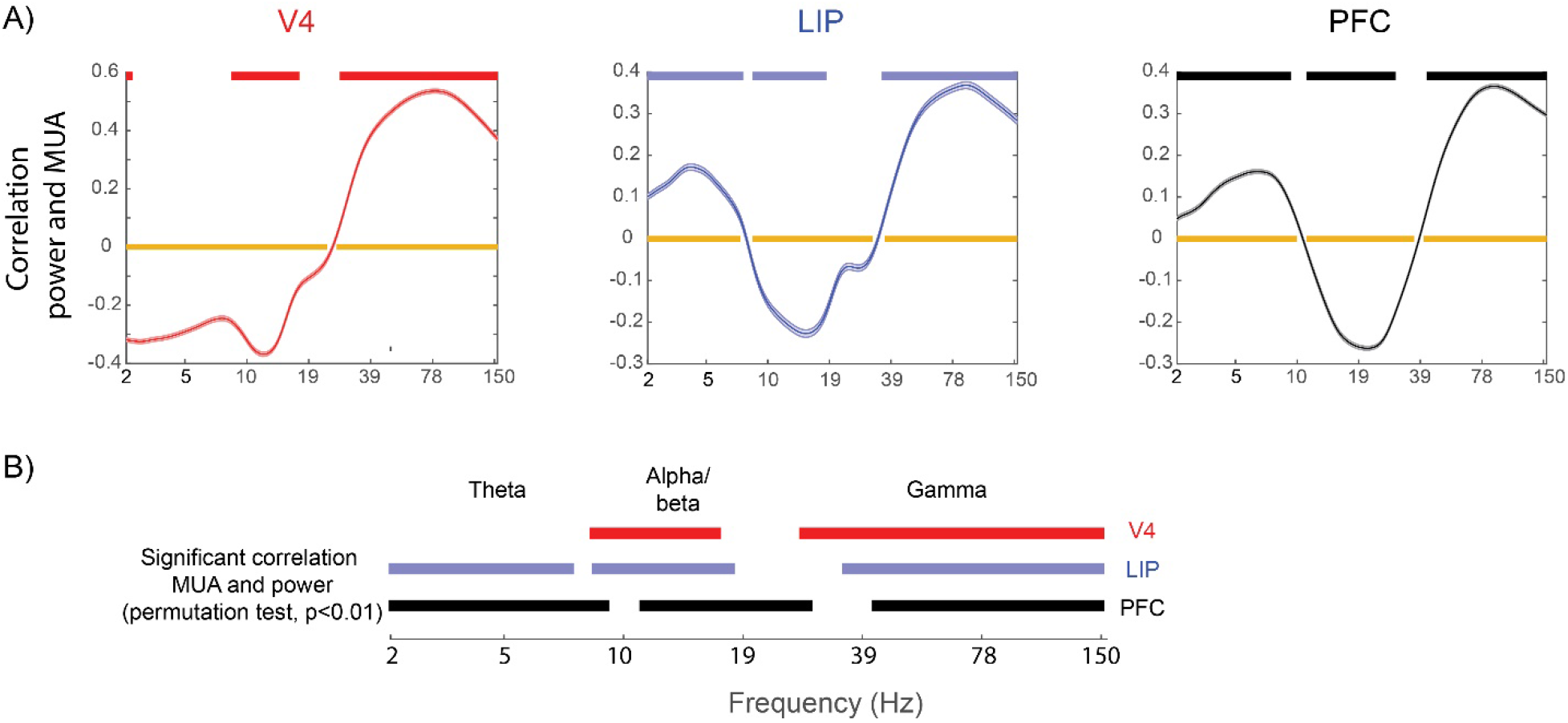
Correlation between power and multi-unit activity. A) The correlation (over time, including 500 ms before stimulus onset until end of the delay) between power in each frequency band and multi-unit activity. Orange bars mark frequencies with significant correlation (p<0.01, t-test for non-zero mean, bon-ferroni corrected). Red/blue/black bars mark frequencies with significantly stronger correlation between power and MUA for neighboring electrodes compared to randomized pairs (permutation test, p<0.01). This was used to control for shared task epoch correlates. B) The red/blue/black bars from A) showing significant correlation for easier comparison.

The peak frequency of maximal negative correlation between LFP power in the alpha/beta band and MUA gradually shifted towards higher frequencies from V4 to LIP to PFC (Figure 3A). Likewise, we defined the frequency of transition between gamma and beta as the lowest frequency above 15 Hz with significant positive correlation with spiking (see Methods). This also increased in frequency up the cortical hierarchy from V4 (26 Hz), LIP (33 Hz), PFC (39 Hz) (all means significantly different at p<10^−15^ using unpaired t-test, corrected for multiple tests). As a result of the changes in frequencies, the frequencies of (positively correlated) gamma in V4 overlapped with (negatively correlated) beta in PFC and PFC (positively correlated) theta overlapped with V4 (negatively correlated) alpha/beta (Figure 3B). Note that the correlation between spiking and gamma did not increase monotonically with frequency. Instead, it peaked below the maximum frequency tested (150 Hz). This suggests that the positive correlations were not due to spikes bleeding into the power spectrum (along with the fact that we used only spikes from neighboring electrodes). Similar results were obtained when we correlated stimulus information in MUA (instead of spike rate) and power (Figure S1).

The correlation between changes in MUA and oscillatory power was not indirect, a by-product of both changing in response to external events (e.g., stimulus onset or offset). Rather, the correlation was direct. To show this, we performed the same analysis but using MUAs and LFPs from random pairs of electrodes from each area instead of neighbouring pairs. Any correlation between such random pairs was attributed to correlation due to them both responding to external events, not a direct correlation. To determine direct correlation, we computed when the average correlation between MUA and LFPs in neighbouring electrode pairs exceeded that of the average correlation between random pairs (permuted 1,000 times). This revealed significant direct correlations (p<0.01), albeit more narrow-band in the alpha/beta domain (Figure 3A, red/blue/black bars). The negative correlations between LFP and spiking were now seen around the peak frequency of the alpha/beta power for each region while gamma remained broadly correlated with spiking. Overall, the patterns of correlations suggested that functionally similar frequency bands (excitatory gamma and theta and inhibitory alpha/beta) in all areas gradually shifted towards higher frequencies up the cortical hierarchy (Figure 3A; but no theta in V4).

We also compared, between electrodes, the correlations between the change in LFP power and the change in MUA from fixation to the memory delay period (Figure 4, using Spearman’s ranked correlation. Bars show significance at p<0.01, Bonferroni corrected for multiple comparisons). This showed that the recording sites with the largest decreases in alpha/beta power during the memory delay also had the highest increases in spiking during that time. In this analysis, theta was positively correlated with spiking in V4 (unlike the other analyses, see above). A similar pattern was seen if the correlation was performed between change in power and change in spiking from baseline to sample stimulus presentation (instead of change from fixation to delay Figure S2).

**Figure 4.**
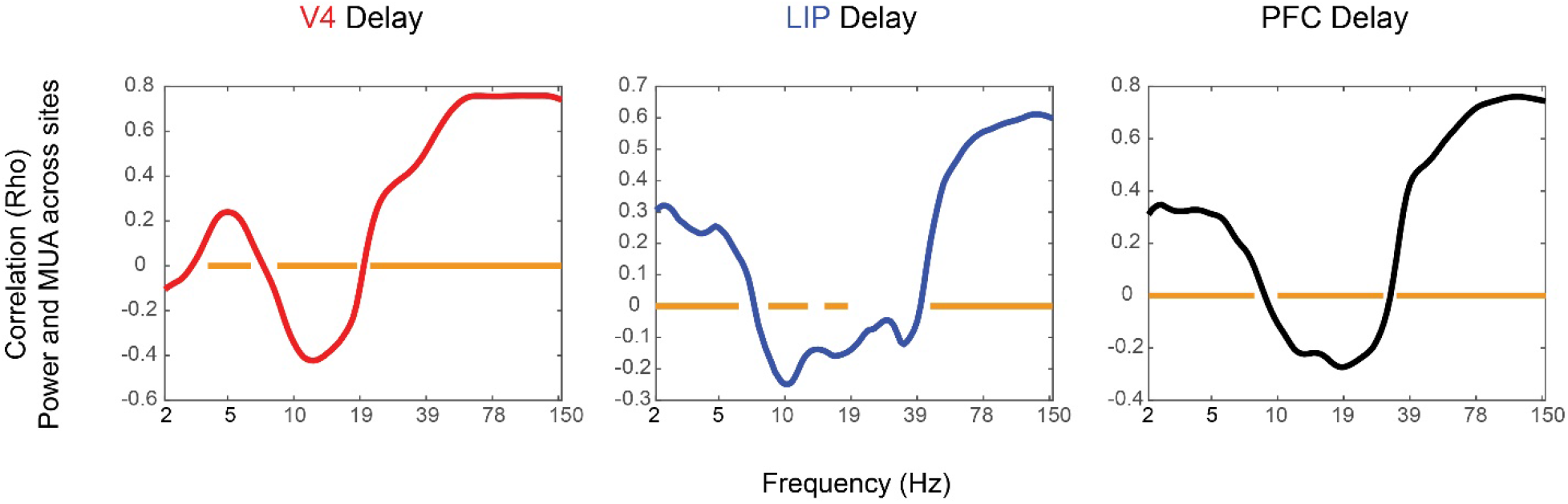
Power and MUA correlation during delay period. Ranked correlation across sites between change (using last 500 ms of fixation and the first 700 ms of working memory delay) in spiking activity (MUA) and change in power. Orange bars show significant correlations at p<0.05, Bonferroni corrected for multiple comparisons.

### Differences in delay activity across the cortical hierarchy

During sample presentation, alpha/beta power was suppressed relative to baseline in all areas but towards the end of the presentation less in V4 than in LIP and PFC (Figure 5A, using 12-26 Hz for all electrodes as this was the alpha/beta range shared by all areas). During the memory delay, alpha/beta power was highest in V4, second highest in LIP, and lowest in PFC (all areas significantly different at each time point at p<0.01, corrected for multiple comparisons, using non-paired t-test). Gamma power during sample presention was highest in V4 (which similar to the suppression in alpha/beta showed a sharper transient to sample presentation), second highest in LIP and lowest in PFC (Figure 5B). During the memory delay, gamma power level was similar across all areas early in the memory delay (Figure 5C) but PFC and LIP showed a ramping up of gamma power towards the end of the memory delay (Figure 5C) while V4 showed a ramping down of gamma power. At the same time, alpha/beta power ramped up in V4 and ramped down in LIP and PFC (Figure 5A). Spiking activity showed similar dynamics and differences across areas as gamma power (Figure 5D).

**Figure 5.**
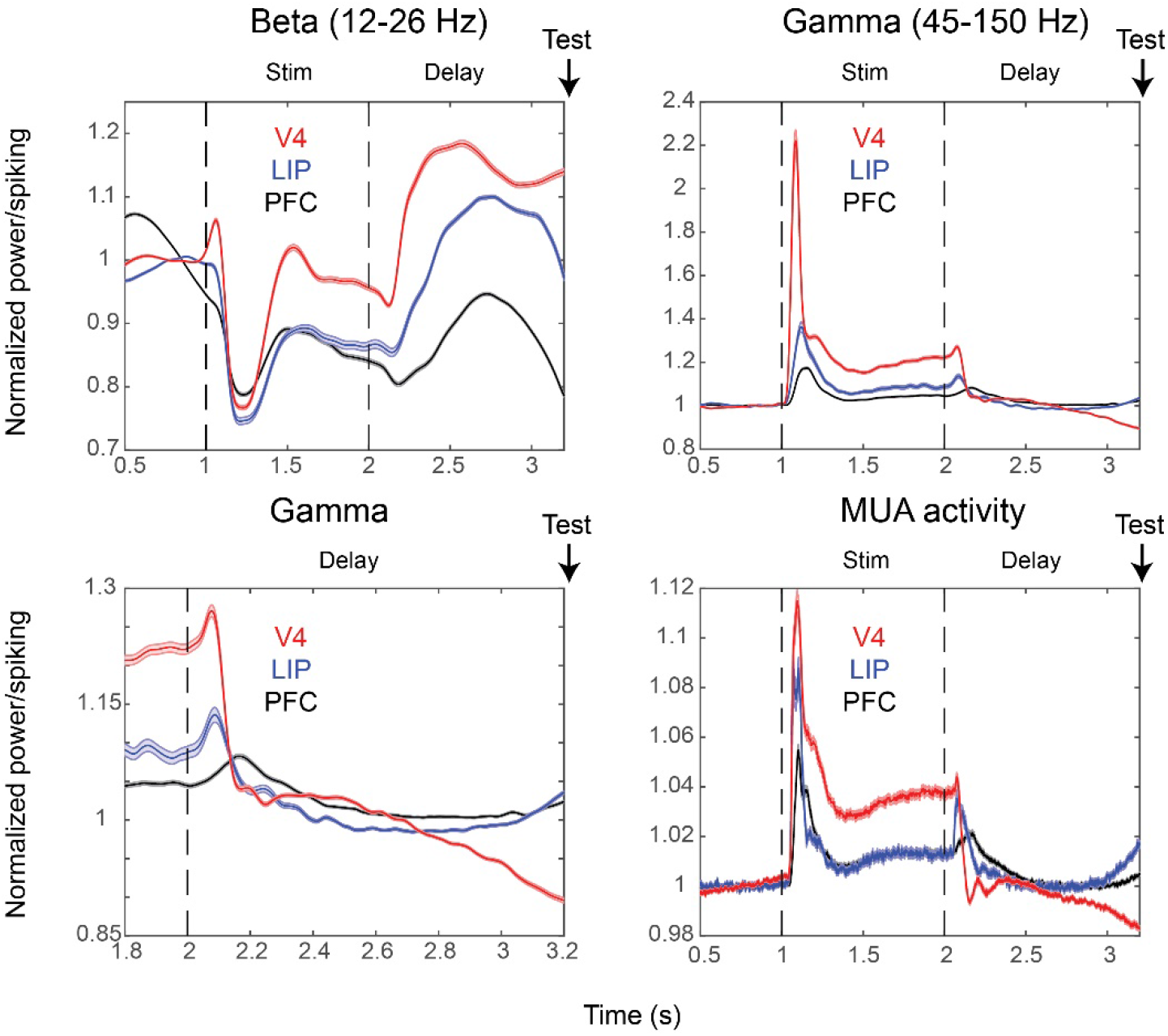
Activity profiles over time. The amplitude of beta (10-27 Hz, A), gamma (45-120 Hz, B, C) and MUA (D) over time, normalized by baseline (0-500 ms before stimulus onset. V4 (red, n=948), LIP (blue, n=528) and PFC (black, n=1560) are shown. Only recordings from sessions in which the delay was held fixed at 1.2 s are used to average signals with similar time-evolution and to display the anticipatory ramp-up. Shaded areas show standard error of mean.

### Gamma bursts were briefer in higher cortical areas

An analysis of single-trial (rather than trial-averaged data) showed that increases in gamma and alpha/beta power occurred in brief bursts. Figure S3 shows example electrodes from each area, demonstrating that the peak frequency differences in alpha/beta were driven by brief high power events around those peaks. To quantify this, we estimated, on each trial, the duration of the bursts and the central frequency at which they occurred. We did so by first setting a threshold at 2 standard deviations above the mean power for each frequency. Around the events where power exceeded the treshold for at least one oscillatory cycle, we fitted 2-D gaussians to estimate their frequency and time (Lundqvist et al., 2016; Methods).

This revealed alpha/beta and gamma occurred in brief and narrow-band bursts in all areas (Figure 6; Figure S3, Methods). Figure 6A shows the duration of the bursts of the stimulus-evoked gamma (i.e. bursts with their central frequency within +−10 Hz of the peak of stimulus-induced power, see Methods and Figure 2). The burst durations were significantly shorter in PFC (mean=37 ms, n=28403) vs LIP (mean=42 ms, n=16177), in PFC vs V4 (mean=47 ms, n=89911) and in LIP vs V4, p<10^−15^ for all comparisons, Wilcoxon rank sum test for equal medians). This was partly, but not entirely, a consequence of the changes in gamma frequency peak across cortical areas. The duration of the bursts inversely scaled with their frequency (combining bursts from all areas; r=−0.52 for beta, p<10^−30^; r=−0.21 for gamma, p<10^−30^). However, selecting bursts from the same frequency range across areas (60 to 80 Hz) resulted in average gamma burst durations that were more similar but still significantly different across areas (Figure S4; Wilcoxon ranked sum test p<10^−09^). Note, for example, that bursts with a central frequency of 26 Hz, which was at the transition to gamma band in V4, had a longer duration (mean 92 ms) than at around the peak. Beta bursts, by contrast, even though their peak frequencies increased along the cortical hierarchy, did not have a corresponding change in duration. Instead they had longer duration in LIP, next longest in V4 and shortest in PFC (Figure 6; LIP mean=240 ms) vs V4 (mean=220 ms), LIP vs PFC (mean = 160 ms), V4 vs PFC, all significantly different medians using Wilcoxon rank sum test with p<10^−15^).

**Figure 6.**
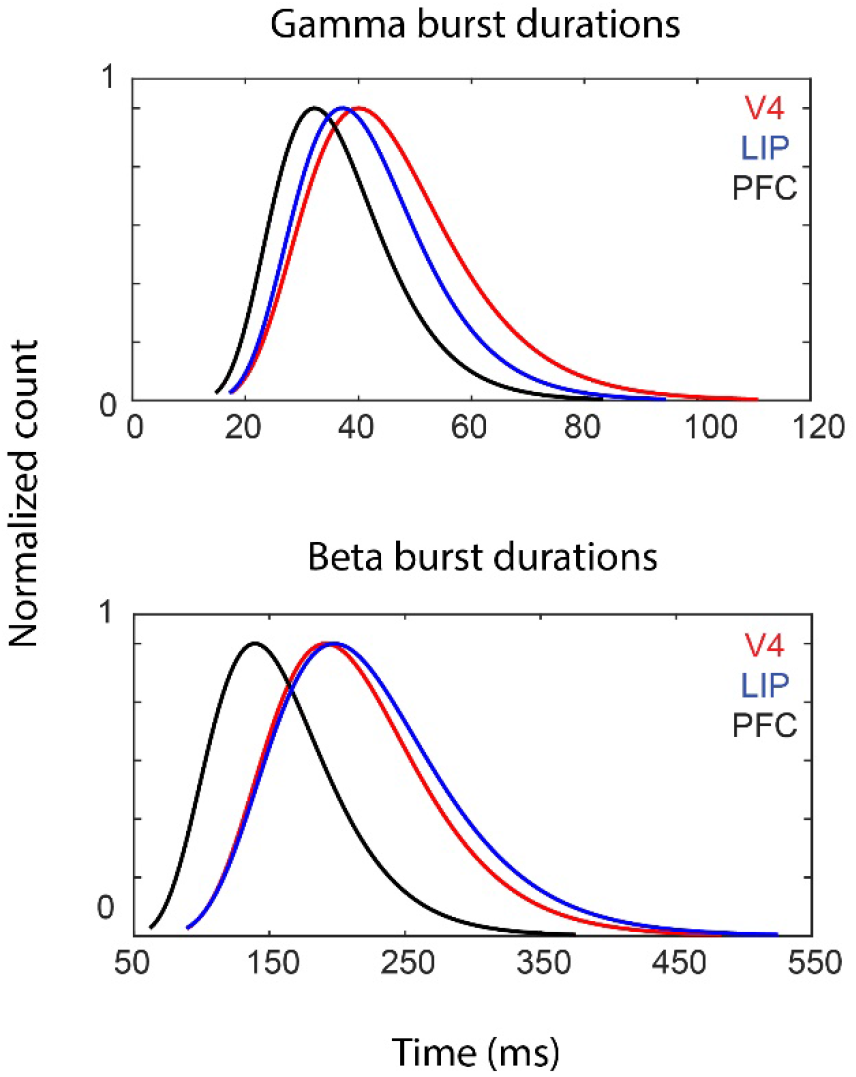
Burst durations per area. Plots show the lognormal fits to the distribution of burst durations for gamma (A) and beta (B), for V4 (red), LIP (blue) and PFC (black). Curves are normalized with respect to maximal count.

In the alpha/beta band there was a clear baseline peak in all areas (Figure 1). Comparisons of all bursts across a wide range (6-35 Hz, using fixation time period) suggested that the differences in average peak power between areas could be explained by differences in central frequency of bursts (p<10^−30^ for all comparisons, Wilcoxon rank sum test for equal median central burst frequency; Figure S3). Potentially, differences in peak power could be due to distinct, non-sinusoidal wave-shapes across compared areas. The power estimate of non-sinusoidal wave shapes contributes power also at higher frequencies than that of the principal period of the wave shape. If this occured more so in for instance PFC it would shift the estimated peak towards higher frequencies despite having the same underlying frequency of oscillation. To rule this out, we used the bottom (rather than central) frequency of each burst. This confirmed the gradually higher alpha/beta frequency up the hierarchy (p<10^−30^ for all pair-wise comparisons, Wilcoxon rank sum test for equal medians).

## Discussion

We observed a general motif as well as a trend across the cortical hierarchy. High frequency (gamma band) power correlated positively with spiking as did slower frequency (theta band) power. By contrast, alpha/beta band power correlated negatively with spiking. The peak frequency of all of these frequency bands became systematically higher as the cortical hierarchy was ascended from visual to parietal to prefrontal cortex. Note that because the peak frequencies of all frequency bands gradually increased up the cortical hierarchy, there was some overlap between “excitatory” (positive correlation with spiking) and “inhibitory” (negative correlation with spiking) frequencies across areas. For example, the excitatory theta in PFC overlapped with the inhibitory alpha/beta in V4. The fast excitatory rhythm (gamma) of V4 overlapped with the inhibitory rhythm (alpha/beta) of PFC. This suggests that the origin of the rhythms should be taken into account when determining the functional relevance of different frequency bands. The power increases occurred in bursts. The duration of the gamma bursts was shorter in higher areas. Finally, in the higher areas (PFC, LIP) there was a ramping up of gamma bursting (and associated spiking) towards the end of the memory delay (V4 instead ramped down). The ramping up has been linked to the read-out of information from working memory (Hussar and Pasternak, 2010; Lundqvist et al., 2016; 2018).

The shorter duration of the gamma bursts higher in the hierarchy was partly due to the increase in gamma frequencies but it may also be a consequence of increases of frequencies in the alpha/beta band. In posterior cortex, gamma bursts are strongly coupled to the ongoing phase of alpha (Osipova et al., 2008; Spaak et al., 2012; Voytek et al., 2010). In frontal cortex, there is phase amplitude coupling between beta and gamma (Bastos et al., 2018). This difference can be explained by the increases in alpha/beta frequencies from posterior to anterior cortex. Gamma will naturally entrain to whatever frequency is most prominent in a given area. The higher frequency alpha/beta peak frequency in PFC may therefore partly explain the shorter gamma burst durations in PFC. The length of the average “duty cycle” (the depolarized or active phase of an oscillation) of a beta frequency is shorter than the average duty cycle length of an alpha oscillation (Figure 7).

**Figure 7.**
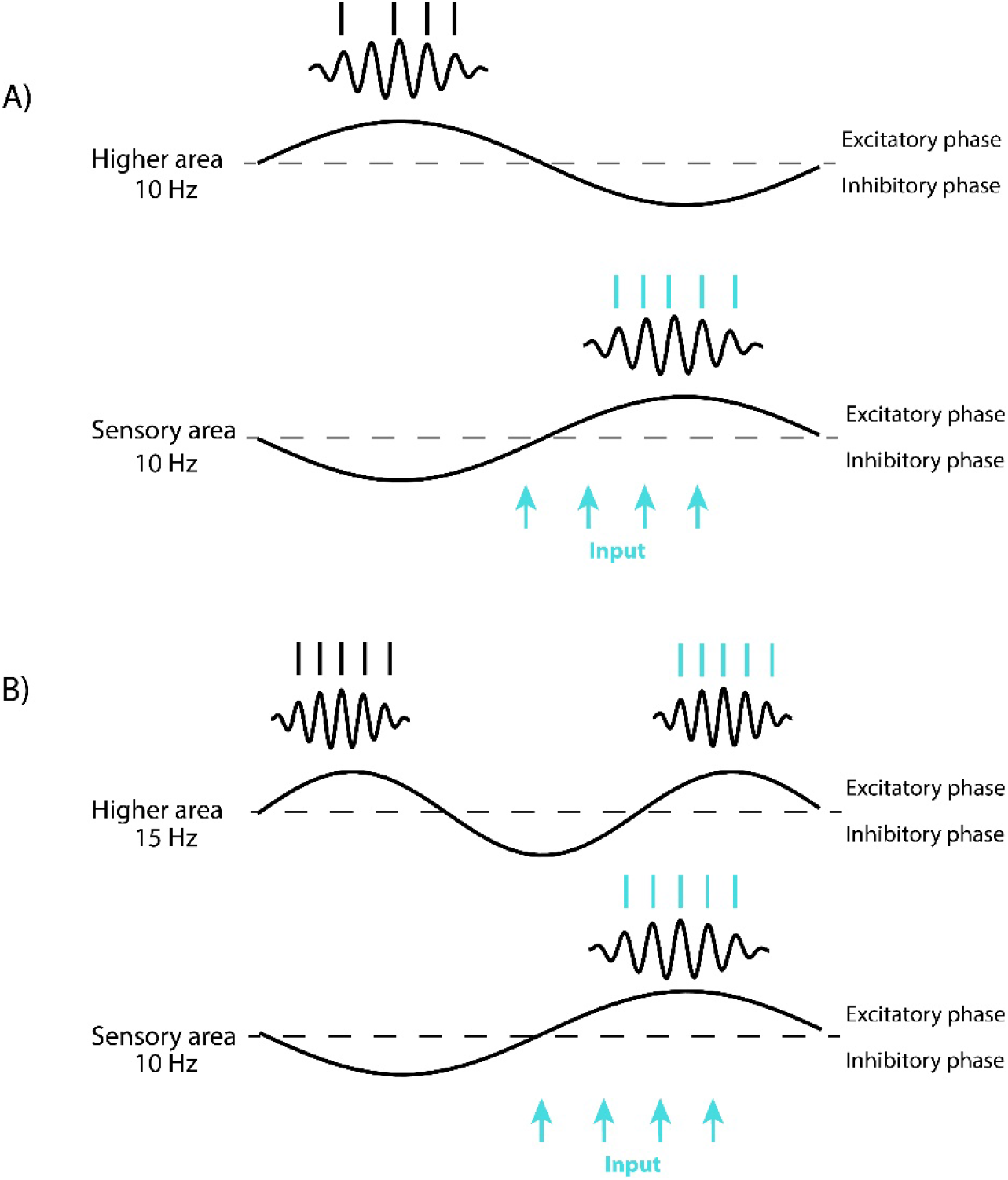
A consequences of increases in alpha/beta frequencues up the cortical hierarchy. Dotted lines denote spike thresholds, above which there is spiking and bursts of gamma oscillations. This active “duty” cycle becomes shorter if the underlying frequency is increased resulting in shorter windows of activity. In the plot, the higher area and sensory area are maximally phase shifted. The blue arrows represent a sensory input. If both sensory and higher areas are sampling at the same frequency (A), a sensory signal will be missed if the areas are out of phase. If the higher area is sampling at a higher frequency (B), the duty cycles will always partly overlap, thus fostering a higher area’s ability to sample sensory inputs. Thus, while alpha/beta can have an overall suppressive effect on gamma/spiking by introducing windows of inhibition, the increase in alpha/beta frequencies in higher areas can help feed forward what spiking there is.

Alpha/beta and gamma rhythms are common and anti-correlated across the cortex but it has been unclear whether or not they have similar functional relationship in different areas. Frontal beta rhythms have been associated with top-down attention, inhibiting motor actions, and working memory storage (Schmidt et al., 2019: Lundqvist et al., 2016; 2018; Buschman and Miller, 2007). Occipital and parietal alpha has been linked to inhibition of unattended stimuli (Klimesch et al., 1998; Worden et al., 2000; Jokisch and Jensen, 2007; van Ede et al., 2011; Bollimunta et al., 2011; Rohenkohl and Nobre, 2011). Gamma power increases (and alpha/beta power decreases) in sensory cortex during sensory input and before/during movements in motor cortex (Brovelli et al., 2004; Cheyne et al., 2008; Schmidt et al., 2019; Fries et al., 2001, Gray et al., 1990; Gevins et al., 1997; Fisch et al., 2009). Frontal beta, and premotor alpha are correlated with reduced spike rates (Haegens et al., 2011; Lundqvist et al., 2016; 2018). Our results suggest that these alpha to beta frequency differences are part of a continuum across the cortical hierarchy with similar general functions. Indeed, regardless of the exact frequencies of alpha, beta, or gamma, we found similar functional relationships at each cortical level. Alpha/beta and gamma were negatively and positively correlated with spiking, respectively, and negatively correlated with each other. Both alpha/beta and gamma were similarly modulated by stimulus processing and the WM task across the entire hierarchy. On a finer scale, we also observed differences with more alpha/beta during WM retention in the sensory areas, consistent with the proposed inhibitory function and more sustained delay activity in higher order cortex. Theta was not as widely observed across cortex but in LIP and PFC it, like gamma, was positively correlated with spiking. In V4, a peak in theta range, positively correlated with spiking during the memory delay.

Modelling suggests that increased peak frequency of oscillatory rhythms in higher cortical areas can arise naturally due to inputs from earlier areas contributing to increased excitation in the higher-order areas (Lundqvist et al., 2013; Brunel and Wang, 2003). Increases in excitation up the cortical hierarchy has also been inferred from large scale connectivity data (Chaudhuri et al., 2015). At the same time, the auto-correlation of spiking suggets a slower time constant for spiking in higher than sensory cortex (Murray et al., 2014). This is not incompatible with our observations of higher, not slower, oscillatory dynamics in higher-order cortex. Rather, spiking vs oscillatory LFPs rhythms could reflect different mechanisms with different functions. The slower dynamics for spiking can help higher-order cortex integrate information from lower areas (Chaudhuri et al., 2015). By contrast, the increase frequency in oscillatory rhythms may help sculpt information flow in cortex. Below, we elaborate.

While there may be a shared inhibitory control function for alpha/beta rhythms across cortex, increases in oscillatory frequencies can still be functionally relevant (Bosman et al., 2012; Wutz et al., 2018). Spiking favoring specific oscillatory phases can group spiking into ‘packets’ of information (Buschman and Miller, 2010; Schroeder and Lakatos, 2009). According to the ‘inhibition-timing’ hypothesis, alpha oscillations reflect alternating periods of relative inhibition and excitability (Klimesch et al., 2007; Haegens et al., 2011). The frequency of the oscillation dictates the duty cycle and thus the duration of nested gamma/spiking activity during the excitatory phases of the alpha rhythm. Different inputs arriving during the same duty cycle of a cortical alpha rhythm tend to be integrated (Van Rullen and Koch, 2003). For example, the temporal resolution at which humans can discriminate two flashes in close succession is finer in subjects with higher occipital alpha peak frequency (Samaha and Postle, 2015; Wutz et al., 2018). This is because the higher the occipital alpha frequency the more likely that the two flashes will arrive at different duty cycles and therefore be perceptually segregated. Further, a recent MEG study showed that the alpha/beta rhythms are not fixed but can flexibly change with changes in task demands for perceptual segregation (Wutz et al., 2018). Alpha peak frequency increased when two successive flashes needed to be segregated and decreased when integration was demanded. Such flexible control over the speed of ‘alpha/beta clocks’ might be achieved by adjusting top-down drive from areas higher in the hierarchy. Increased top-down drive would entrain the sensory alpha to a slightly higher frequency.

In the context of a hierarchy of connected areas, it also makes sense that the ‘alpha/beta clocks’ increase in frequency up the cortical hierarchy. It allows higher areas to better segregate packets received from lower cortical areas and reduces the risk that two packets will be merged into one. Further, while increases in alpha/beta *power* can suppress gamma/spiking by introducing periods of inhibition (Haegens et al., 2011), the increases in alpha/beta *frequencies* up the cortical hierarchy can help feed forward whatever spiking there is. If the receivers higher in the cortical hierarchy operate on a slightly higher time scale, it will make it more likely that the receiver will have some overlap between its duty cycle and incoming packets sent during the duty cycle of the sender, supporting the feedforward flow of spiking (Figure 7).

At the same time, the increase in frequences up the cortical hierarchy may also bias the direction of rhythmic entrainment in the cortex in the feedback direction. If two oscillators with different harmonic frequencies are connected, the higher oscillator will be more effective at entraining the slightly slower oscillator, than vice-verse. The increase in frequencies in higher areas would thus naturally bias rhythmic entrainment in the feedback direction, down the cortical hierarchy. When there are no sensory inputs, alpha/beta rhythms predominate in cortex. They have been associated with top-down control (Buschman and Miller, 2007; Bastos et al., 2015; Van Kerkoele et al., 2014). The frequency gradient we observed suggests that their entrainment has a feedback directional bias consistent with top-down control. Bottom-up inputs could overcome the top-down bias during periods of strong sensory drive. Since sensory stimuli drive gamma, it could explain the entrainment of gamma rhythms predominantly in the feedforward direction. In sum, the increase of oscillatory frequencies up the cortical hierarchy may provide another mechanism by which cortical processing and flow can be regulated.

## Acknowledgements

We would like to thank Pawel Herman and Nancy Kopell for useful discussions. This work was supported by National Institutes of Mental Health Grant R37MH087027 and 5K99MH116100-02, Office of Naval Research Multidisciplinary University Research Initiatives Grant N00014-16-1-2832, the MIT Picower Institute Faculty Innovation Fund, Swedish Research Council Starting grant 2018-04197 and the 2017 Young Investigator Grant from the Brain & Behavior Research Foundation.

---

**Supplementary Figure S1.**
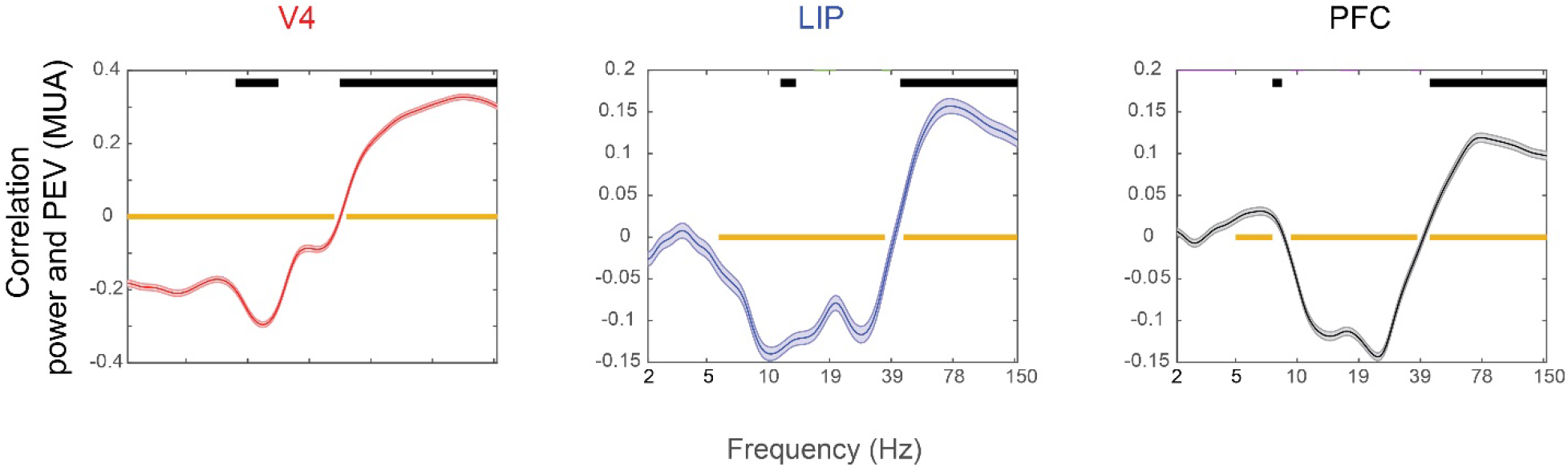
Correlation between power and multi-unit information. A) The correlation (over time, including 500 ms before stimulus onset until end of the delay) between power in each frequency band and percentage explained variance (PEV) by object sample cue identity in MUA. Orange bars mark frequencies with significant correlation (p<0.01, t-test for non-zero mean, bon-ferroni corrected). Black bars mark frequencies with significantly stronger correlation between power and MUA for neighboring electrodes compared to randomized pairs (permutation test, p<0.01). This was used to control for shared task epoch correlates. Shaded areas show standard error of mean.

**Supplementary Figure S2.**
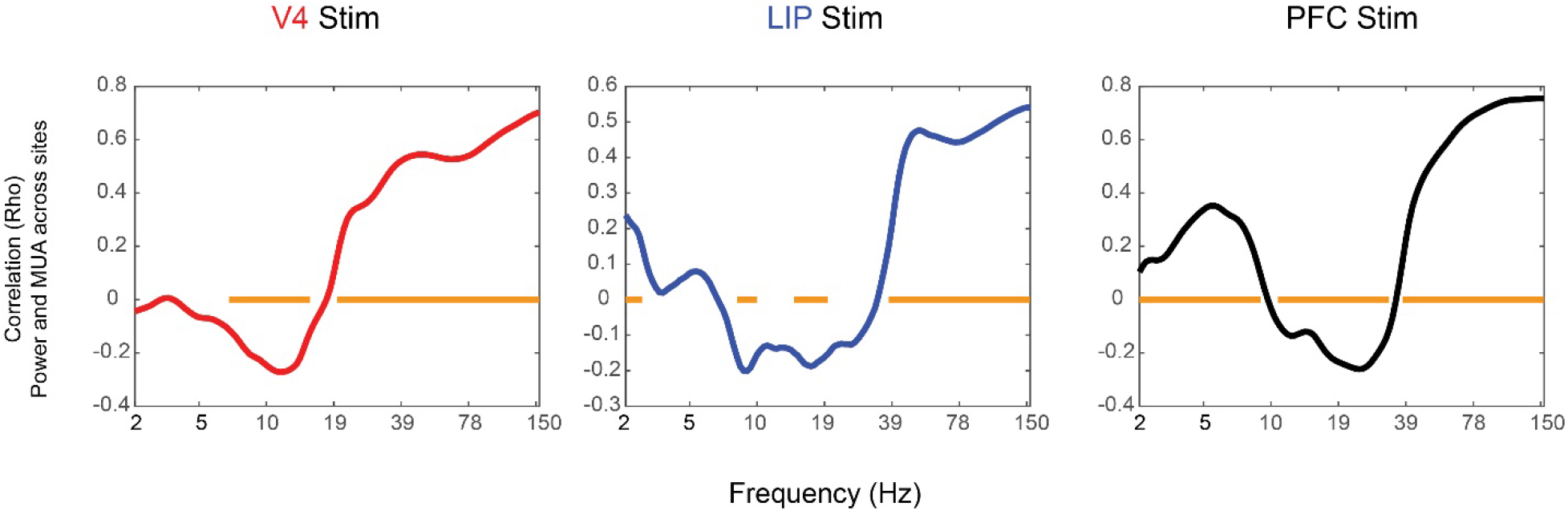
Power and MUA correlation during stimulus. Ranked correlation across sites between change (from fixation to stimulus) in spiking activity (MUA) and change in power. Orange bars show significant correlations at p<0.05, corrected for multiple comparisons.

**Supplementary Figure S3.**
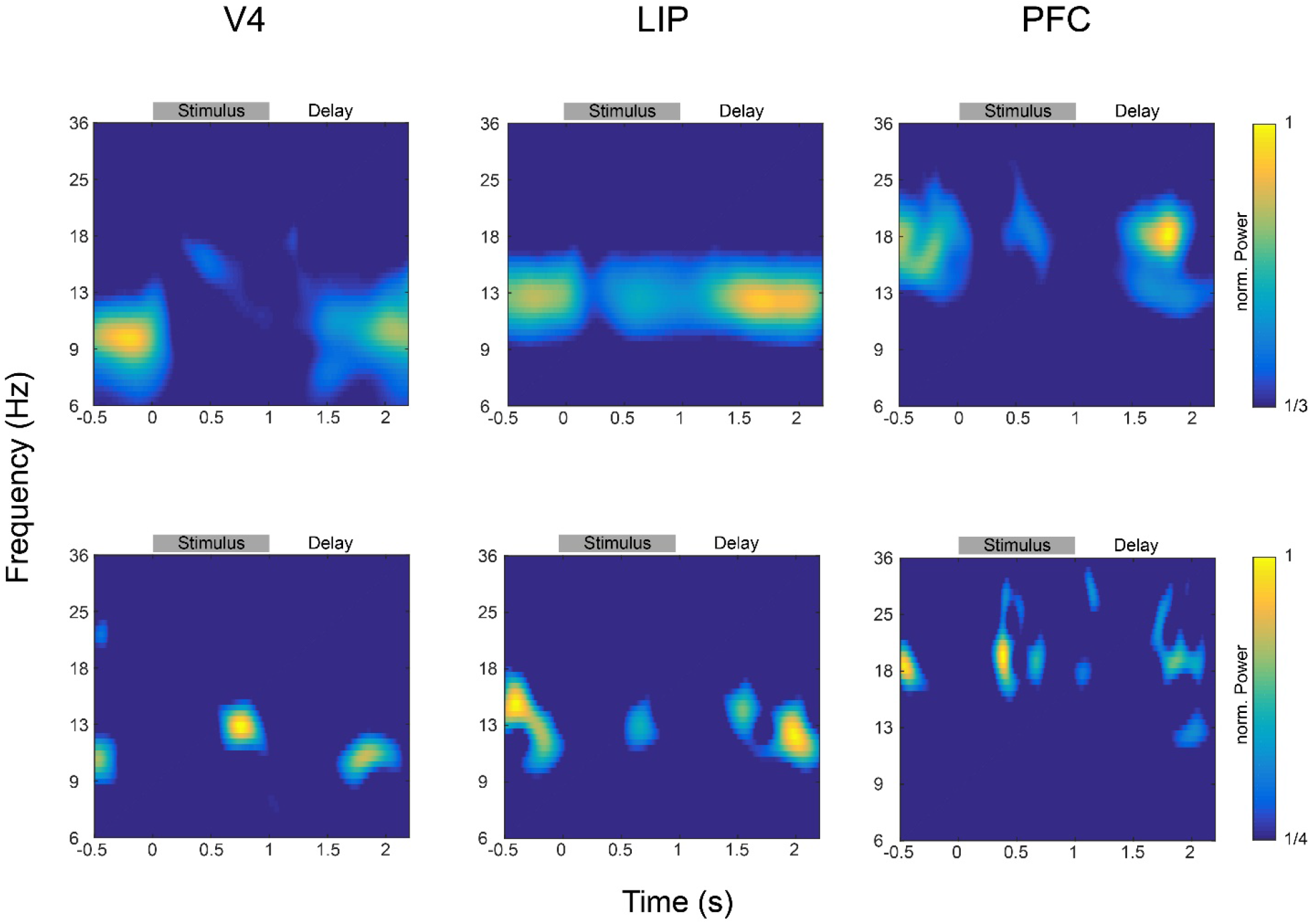
Alpha/beta activity in example electrodes. Representative V4, LIP and PFC example electrodes recorded during the same session showing time-frequency representations in the alpha/beta range. Top row shows the trial-averaged power for all correct trials, bottom row shows single trial examples. Power is normalized to the maximum value in of each plot.

**Supplementary Figure S4.**
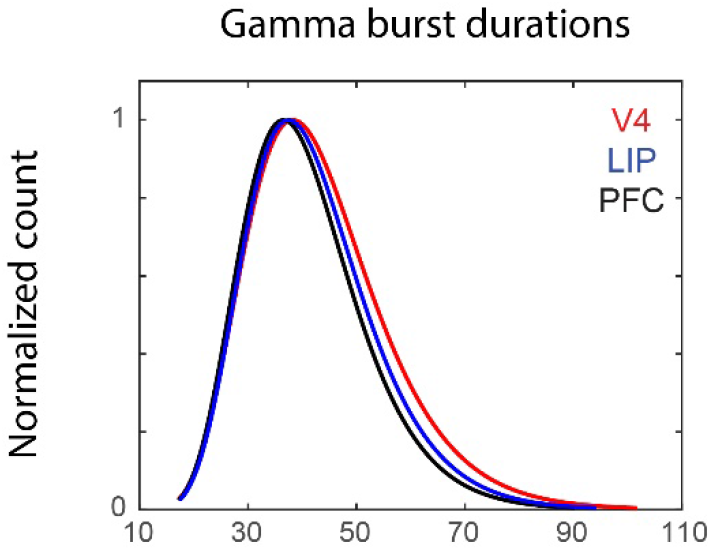
Gamma burst durations per area. Plots show the lognormal fits to the distribution of burst durations for V4 (red), LIP (blue) and PFC (black) between 60 and 80 Hz. Curves are normalized with respect to maximal count.

