## Supplemental Figures 1-3 for "Preservation and changes in oscillatory dynamics across the cortical hierarchy"

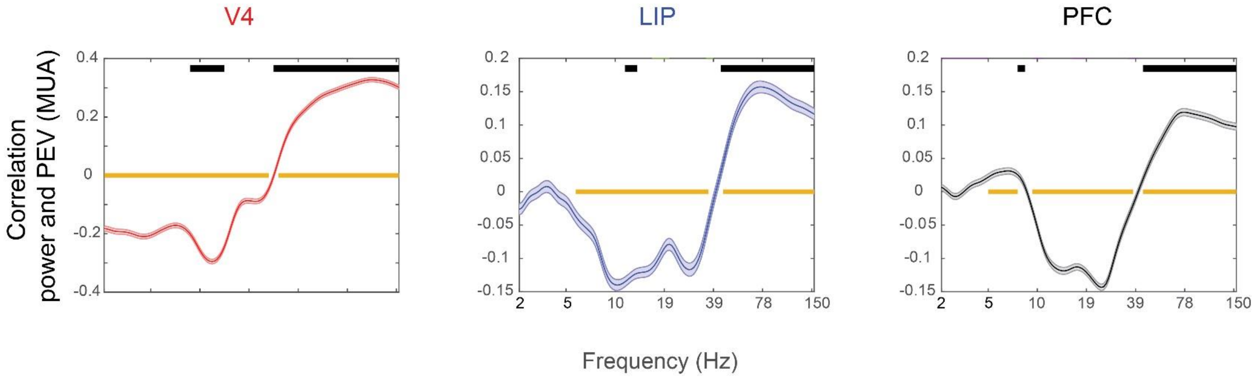


Supplementary Figure S1. Correlation between power and multi-unit information. A) The correlation (over time, including 500 ms before stimulus onset until end of the delay) between power in each frequency band and percentage explained variance (PEV) by object sample cue identity in MUA. Orange bars mark frequencies with significant correlation (p<0.01, t-test for non-zero mean, bon-ferroni corrected). Black bars mark frequencies with significantly stronger correlation between power and MUA for neighboring electrodes compared to randomized pairs (permutation test, p<0.01). This was used to control for shared task epoch correlates. Shaded areas show standard error of mean.

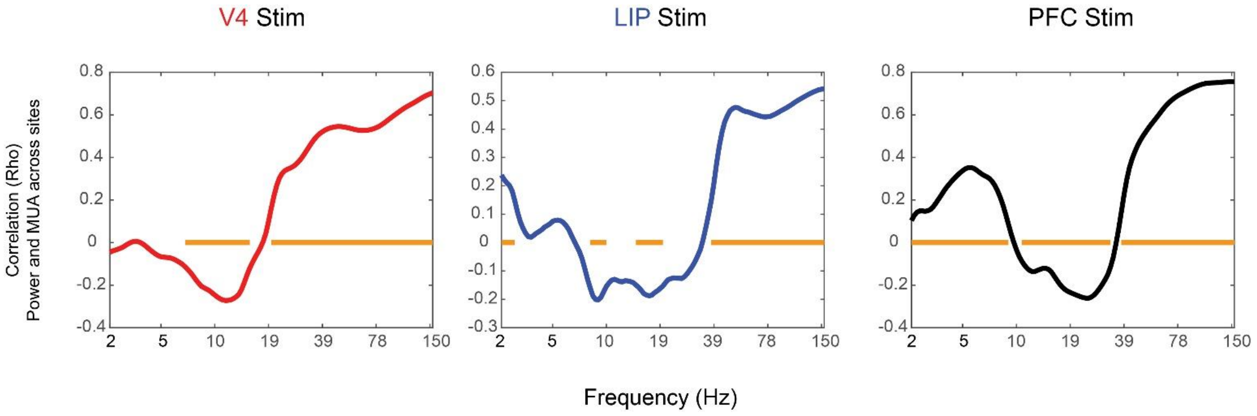


Supplementary Figure S2. Power and MUA correlation during stimulus. Ranked correlation across sites between change (from fixation to stimulus) in spiking activity (MUA) and change in power. Orange bars show significant correlations at p<0.05, corrected for multiple comparisons.

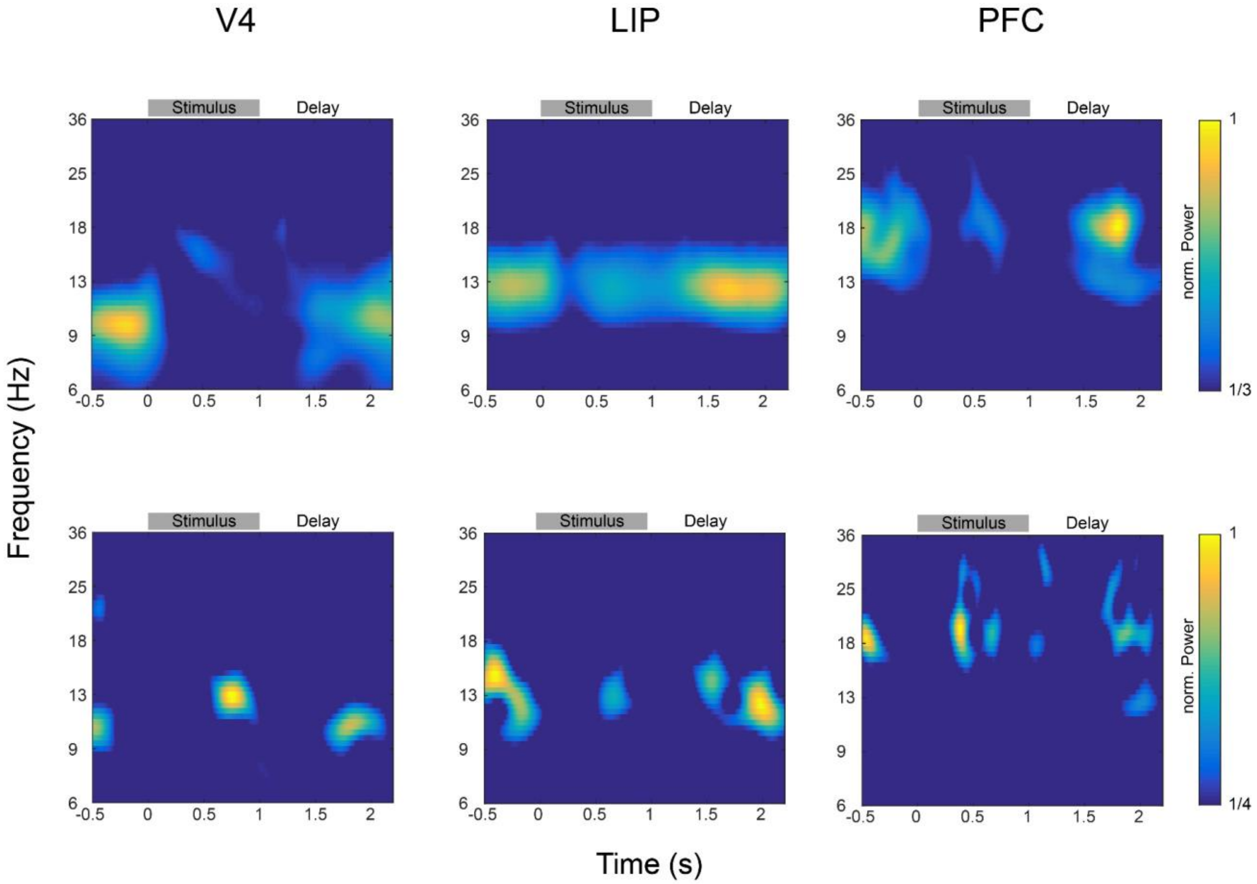


Supplementary Figure S3. Alpha/beta activity in example electrodes. Representative V4, LIP and PFC example electrodes recorded during the same session showing time-frequency representations in the alpha/beta range. Top row shows the trial-averaged power for all correct trials, bottom row shows single trial examples. Power is normalized to the maximum value in of each plot.

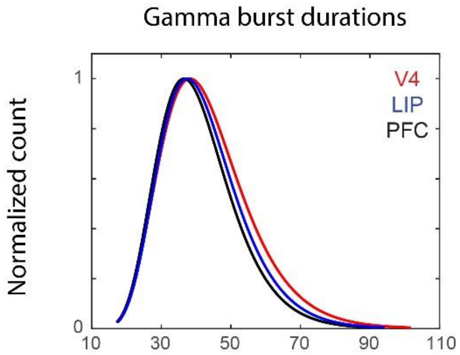


Supplementary Figure S4. Gamma burst durations per area. Plots show the lognormal fits to the distribution of burst durations for V4 (red), LIP (blue) and PFC (black) between 60 and 80 Hz. Curves are normalized with respect to maximal count.
